# A mixture of generative models strategy helps humans generalize across tasks

**DOI:** 10.1101/2021.02.16.431506

**Authors:** Santiago Herce Castañón, Pedro Cardoso-Leite, Irene Altarelli, C. Shawn Green, Paul Schrater, Daphne Bavelier

## Abstract

What role do generative models play in generalization of learning in humans? Our novel multi-task prediction paradigm—where participants complete four sequence learning tasks, each being a different instance of a common generative family—allows the separate study of *within-task learning* (i.e., finding the solution to each of the tasks), and *across-task learning* (i.e., learning a task differently because of past experiences). The very first responses participants make in each task are not yet affected by within-task learning and thus reflect their priors. Our results show that these priors change across successive tasks, increasingly resembling the underlying generative family. We conceptualize multi-task learning as arising from a mixture-of-generative-models learning strategy, whereby participants simultaneously entertain multiple candidate models which compete against each other to explain the experienced sequences. This framework predicts specific error patterns, as well as a gating mechanism for learning, both of which are observed in the data.

## Introduction

Understanding the mechanisms underlying human learning remain amongst the most important, intriguing and fascinating open challenges for science. Insights about human learning have the potential for large-scale societal impact, for example, by driving progress in artificial learning systems. Humans are exceptional learners. They can learn effectively from very limited experience (e.g., Carey & Bartlett, 1978; Feldman, 1997). They can generalize their learning beyond the specifics of their experience (Jern & Kemp, 2013; Lake et al., 2015; Xu & Tenenbaum, 2007), such as correctly categorizing previously unseen *exemplars* of a category (T. B. Ward, 1994). Interestingly, both the speed with which humans can learn (i.e., learning from few examples) and the ability to generalize (i.e., transferring knowledge across contexts) may indicate that humans approach learning problems in a structured way (Behrens et al., 2018; Lake et al., 2017). In essence this would involve making use of pre-existing knowledge about the world (i.e., “priors”), and then consistently incorporating new pieces of information into a generative model (i.e., a structured and generalizable model of how the world has generated the given experiences).

In a wide variety of fields, including perception and decision-making, an overwhelming amount of evidence shows that humans and other animals rely on internal generative models to make optimal inferences about currently presented information (Behrens et al., 2007; Kersten et al., 2004; Kilner et al., 2007; Körding et al., 2004; Wolpert et al., 1995). A major benefit of using a generative model is that such models allow for inferences and predictions to be made about future data. Accordingly, generative models have also been shown to play a critical role in guiding sequential behaviour (Cleeremans & McClelland, 1991; Elman, 1990), where forecasting of future scenarios, planning and preparation of future actions, and the use of counterfactual reasoning are all crucial for success (Boorman et al., 2009; Daw et al., 2005; Koechlin et al., 1999; G. Ward & Allport, 1997). For instance, after learning to navigate through a maze, internal models of the maze allow individuals to quickly find a short path to a new goal—even when that path had not been previously used (Tolman & Honzik, 1930). Together, these results highlight the important role that generative models play in guiding behaviour. Yet, we know very little about the role generative models play in guiding the learning process itself. We owe this state of affairs, at least partially, to the difficulty of studying the impact of generative models on learning processes. In particular, it is intrinsically the case that those situations where generative models are most useful for guiding learning are also the situations where it is most difficult to study their influence. Below, we provide some of the reasons for this difficulty as well as our approach to circumventing those challenges.

Why, then, is it difficult to study the role of generative models in learning? Quantifying and understanding the role that generative models play during learning has been difficult for several reasons. First, the utility of a generative model in guiding learning is often highest in the earliest phases of learning (i.e., when the learner is still “data poor” and must rely on the generative model rather than direct experience). Yet, because this phase is often short, it is difficult for researchers to track the underlying beliefs that are guiding participants’ learning during this stage (e.g., practically speaking, data cannot be aggregated to effectively reduce measurement noise). Second, it is always the case that other cognitive processes may simultaneously compete to control observable behaviours or choices. For instance, learning processes that do not rely on generative models (e.g., trial-and-error or model-free learning) may be equally or more prominent when behaviour reaches a stable state (i.e., at asymptotic performance), making it difficult to attribute choices to one process over another. Finally, most studies rely on laboratory paradigms where the environment is governed by a single generative process. This is typically the case in temporal sequence learning paradigms, where participants are exposed to sequences of stimuli, often extensively long ones, that are governed by a single generative process. While tasks with a single generative process have been very fruitful in studying how internal generative models guide behaviour (Cleeremans & McClelland, 1991; Knill & Richards, 1996; Pouget et al., 2013; Tenenbaum et al., 2006), it makes it impossible, however, to track (i) how experience in that environment shapes learning in other scenarios, or (ii) what behaviour looks like during learning in an environment that is possibly controlled by competing generative models.

Here we present a novel multi-task paradigm designed to examine the role that generative models play in learning. In this study, participants went through a series of four separate temporal sequence prediction tasks. The sequence of stimuli in each of the tasks was governed by a specific generative process selected from amongst a broader common generative family. Across a series of 10 experiments involving over 800 human participants, we asked two fundamental questions. First, when learning a task, are participants using (or inferring) a generative family? To answer this question, we quantified participants’ prior knowledge as they started each new task. Specifically, the first four trials of each task were purposefully designed to reveal participants’ priors uncontaminated by direct experience with the given task. Thus, by analysing these very early trials we are able to show that participants rapidly and consistently (i.e., starting from exposure to a single instance of the generative family but continuing through the exposure to following instances) update their priors to better match the true generative family when approaching each subsequent task. The second fundamental question we ask is, how do people learn within tasks that are situated in a multi-task generative model environment? We present evidence that participants use a mixture-of-generative-models (MGM) learning strategy. In particular, we see that the MGM learning hypothesis (i) explains specific structure within the errors participants made, (ii) naturally accounts for the large individual variability in behaviour in terms of the prior with which people start the task (as opposed to relying on ad-hoc assumptions to explain suboptimal behaviour), as well as (iii) predicts the existence of a gating mechanism for learning which explains why people with inadequate priors fail to learn the task.

## Results

Participants (*n* = 854, across 10 experiments, see Methods) were placed in a multi-task learning environment in which they were asked to perform a series of sequential prediction learning tasks interleaved with production tasks. A common generative family defined the possible generative models that could control the stimulus sequence in any one prediction task (**Fig.1A**). All possible generative models within the generative family follow a first order Markov process from which sequences of stimulus locations are sampled. The generative family boils down to two pieces of information required to describe any single instance of a generative model from the generative family (see Methods for a full description): (i) all generative models follow one of four dominant transition patterns, i.e., the pattern describing which location is the most likely next location given any current location (see **Fig.1A** top panel), and (ii) one dominant transition probability value, **α**, shared across all locations (i.e., the probability with which the stimulus will appear in the most likely next location). One particular instantiation of this generative family then determines in which of four possible screen locations the stimulus appears on each trial in a given task.

**Fig 1.**
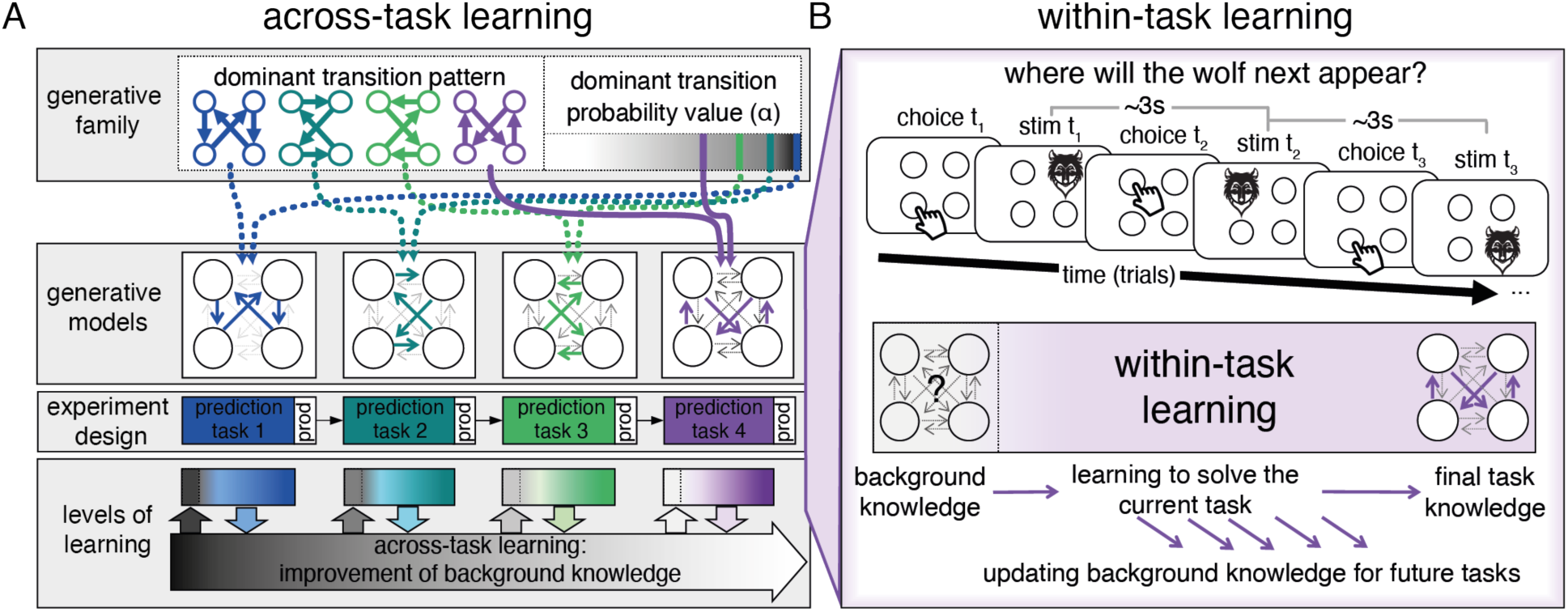
The multi-task learning paradigm. (**A**) ***Across-task learning***: In this multi-task learning environment, participants went through a series of sequential prediction tasks. The stimulus’ probabilistic behaviour in each task is determined by a specific generative model— each generative model is a different instance from a common generative family (top row; also see methods) defined by two characteristics: the identity of the dominant transition pattern (top row, left; note coloured arrows indicate the most likely stimulus transition from each of the four locations on the screen), and the dominant transition probability value, **α** (i.e., the probability of the stimulus moving to the most likely next location; top row, right). These two pieces of information are then combined (second row from top) to create a particular generative model from which the to-be-predicted sequences of locations are drawn (e.g., for prediction task 4, the generative model uses the purple dominant transition pattern along with a small value of **α**). For each generative model, participants first completed a prediction task (where they observed the stimulus appear in one location and then predicted the next) and then a production task (where they were asked to produce a sequence in the absence of any observations; third row from top). Because participants underwent multiple such tasks, learning could occur both within a task as well as across tasks (bottom row). (**B**) ***Within-task learning***: For each prediction task, participants had to guess the next location of “the wolf”. In this, participants were simply told to do their best to “catch the wolf” (i.e., they were given no information about the generative models). Participants had a three-second window to make each prediction (clicking one of the four locations) before the wolf appeared in the next location (for one second). Once the wolf disappeared again, a new prediction could occur (top panel). Critically, the first four choices participants make in each task can only be guided by the background knowledge participants bring to the task, as the wolf does not repeat locations within the those first four trials. Experiencing the wolf transitions in any one task should help participants learn to solve the current task as well as update the background knowledge with which they will approach future tasks (though applying the exact solution of the current task directly to future tasks can be disastrous for performance; bottom panel).

Our multi-task learning paradigm, thus, has an advantage over a single-task learning paradigm in that it allows us to assess both within-task learning (i.e., where participants increase the accuracy of their predictions within a given task) and across-task learning (i.e., where participants generalize what they learn in one task beyond the context of that one task to influence their behaviour on future tasks). For each prediction task, the cartoon face of a wolf appeared successively in one of four locations on a screen following a sequence sampled from a specific generative model (i.e., one location per trial). Participants had to predict the next location of the wolf (**Fig.1B**). For the first few trials of each prediction task, participants had to rely on the background knowledge they brought to the task (i.e., the wolf did not reappear in a previously visited location for the first four predictions; thus, participants could not use previous observations within the task to estimate the most likely next location unless they had intuitive biases about the generative family). Then, as they spent more time in each task, participants had the opportunity to learn the task-specific transition probabilities (i.e., learn the task-specific generative model). Crucially, although the specific generative models that determined the sequences of stimulus locations for each task were drawn from the same generative family, it was nonetheless the case that the exact transition probabilities were very different across tasks (e.g., in the first task, the wolf might move mostly to location 4 when in location 1, while in the second task, the wolf might instead move mostly to location 2 when in location 1). As such, if participants simply reused the previously learned transition probabilities in a new task, they would perform very poorly. However, their experience in a given task could allow participants to change their understanding of how tasks in this environment work—an understanding that might help them learn future tasks.

### Performance reveals within-task learning while early behaviour reveals across-task learning

We first evaluated participants’ performance throughout each of the four tasks by computing the proportion of choices that were made in agreement with the true generative model of that task. Under the true generative model of any one task (i.e., the generative model that produced the actual stimulus sequence of the task), for each trial there is a unique most likely next stimulus location given the current stimulus location. We note that this metric is a better indicator of performance than simple “accuracy” (i.e., tallying the times the wolf was caught), due to the stochasticity of the generative model (which means that even if a choice was in agreement with the true generative model, the wolf may not always have been caught and vice versa).

Examining participants’ performance across the trials of each task reveals, first and foremost, clear evidence of learning. Within each task, participants’ choices across trials, on average, became gradually closer to the performance of an ideal artificial learner agent (**Fig.2A**). Second, the extent to which this is true differs from task to task. In tasks with more stochastic stimulus sequences (i.e., tasks with lower **α** values, such as task 4), learning was slower, and the learning curves saturated at lower values, as compared to tasks with more deterministic sequences. This difference across tasks appears to reflect task difficulty (i.e., performance is “worse” in tasks with more stochasticity). We note that this relationship between stochasticity and learning (i.e., higher stochasticity requires more trials to learn the optimal behaviour) would be true even for an ideal artificial learning agent. However, because the tasks differ in difficulty, it suggests that the metric is not ideal for determining whether participants carried over knowledge from one task to the next.

**Fig 2.**
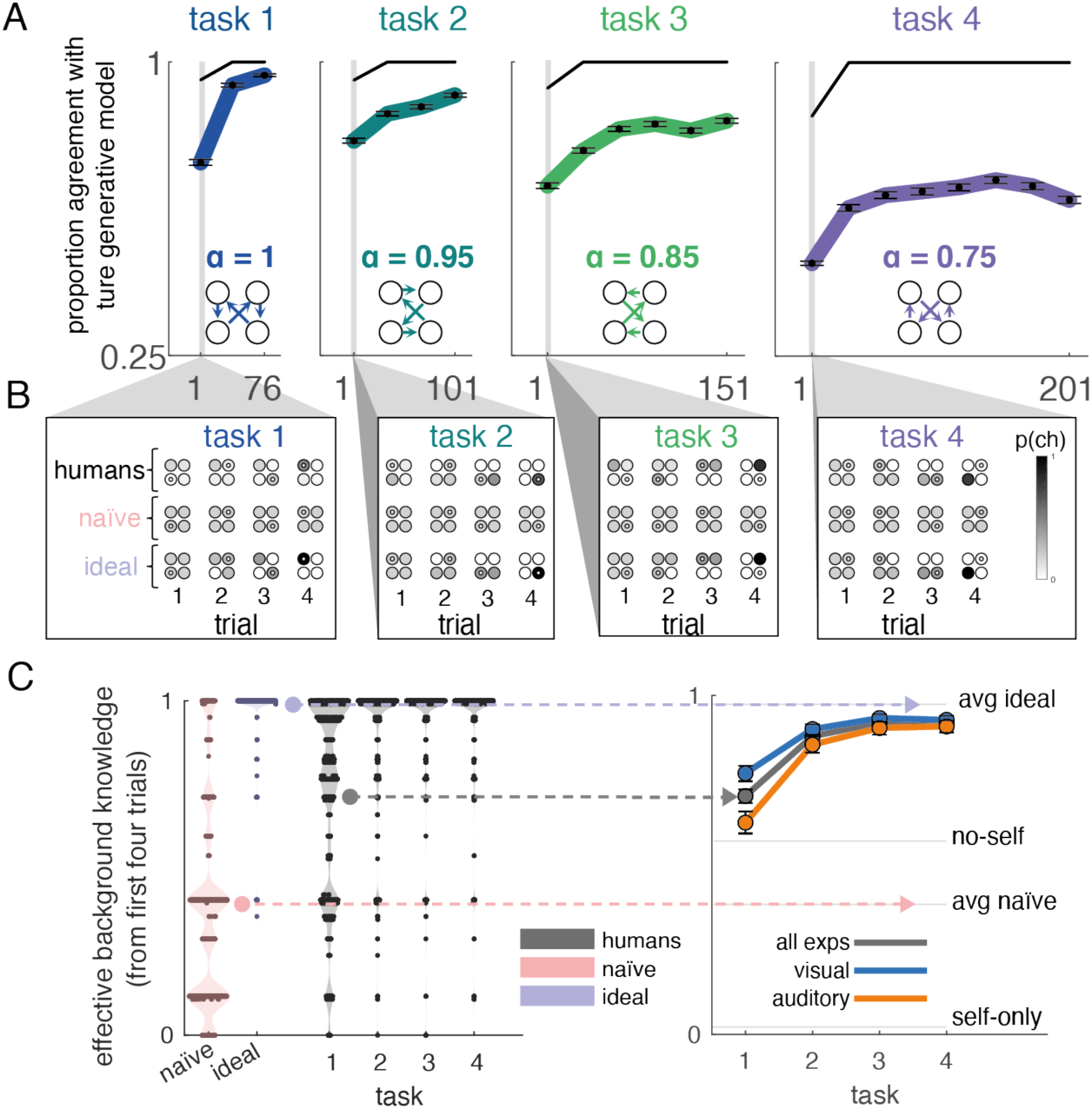
Across-task learning revealed by early behaviour analysis and not by performance. (**A**) *Performance reflects only within-task learning*: Participants (coloured curves, black dots and error bars) rapidly improved their performance in each task (increasing the proportion of predictions in agreement with the true generative model of the task). Even an ideal artificial learner (black upper curve) who knows the generative family, would require a few trials to reach the highest performance level. Task 1 (blue) had the highest dominant transition probability value, **α**, (i.e., it was the least stochastic task) and task 4 (purple) had the lowest **α** (i.e., it was the most stochastic task). As expected, both the speed and asymptotic level of learning were lower for tasks with higher stimulus stochasticity, comparing performance across different tasks is thus not indicative of across-task learning. (**B**) *Early behaviour reveals across-task learning:* the first four predictions of participants have a structured distribution for each of the four tasks (darker colours indicate higher probability; the inner white circle indicates the position the wolf actually appeared on that trial after the choice was made). As reference points, we computed the distribution of predictions under two different artificial learning agents which represent two extremes on a knowledge scale: a naïve artificial learner and an ideal artificial learner (to get comparable distributions, one simulation was done for each artificial learner and for each of our participants). The naïve artificial learner (second row, pink) has no knowledge of the generative family and uses only count statistics to infer the transition probabilities. The ideal artificial learner (third row, blue) has full knowledge of the generative family and uses it to infer the generative model of the current task (e.g., far left panel, bottom row, the ideal agent, after observing the wolf in the bottom left location on trial 1, puts zero probability mass on the wolf appearing in the bottom left location on trial 2, as it knows that such a transition is not possible under the generative family). Already in task 1, participants do not behave like the naïve artificial learner but instead their predictions reveal they arrive with structured intuitions. Furthermore, as participants complete successive tasks, the distribution of their first four predictions increasingly resembles those of the ideal artificial learner, suggesting participants update their background knowledge from one task to another to better match the generative family. (**C**) *Background knowledge improves across tasks.* To establish that participants update their background knowledge from one task to another, we computed the likelihood ratio of participants’ choices (in panel B) relative to the two artificial learners. Distributions represent log-likelihood ratios for human participants and simulated agents rescaled between the extremes of the measure. Participants’ distribution of background knowledge confirmed that already in task 1, participants’ choices (dark grey) deviated strongly from the distribution of the naïve artificial learners (pink) and loosely resembled that of the ideal artificial learners (blue). Critically, background knowledge strongly increased across successive tasks both on average (right side, mean and 95% confidence intervals) and within participants, demonstrating that they increasingly matched their background knowledge to the generative family. This improvement occurred independent of the sensory modality, for both visual (blue) and auditory (orange) versions of the experiment. The apparent discrepancy between overall task performance (panel A) and early task behaviour (panels B and C) is consistent with the mixture-of-generative-models learning framework introduced in the following section.

As noted above, examining across-task learning is challenging. For instance, revealing across-task learning requires differentiating behaviour that is driven by experiences that occurred prior to the current task from behaviour driven by experiences that occurred during the current task. This distinction motivated our approach to look at the very earliest behaviour, specifically from the first four predictions in each of the tasks, in order to characterize across-task learning. Our focus on the first four trials is not arbitrary but is well founded on specific aspects of our experimental design.

Namely, as noted above, the first four choices in each task must be made before a wolf reappears in a previously experienced location for that task and as such, there is no evidence from the current task itself, regarding the probability with which it will transition to any other location. Because of this, the first four choices are a unique window for observing behaviour which is purely driven by the prior background knowledge that the participants brought to the current task and not by evidence acquired from observing transitions within the task. The obvious downside to examining such a small number of choices is that estimates related to any one participant will be noisy, and thus, in order to still get reliable estimates of behaviour, we collected and aggregated data from hundreds of participants.

Focusing on the first four predictions of each task allows us to ask two interrelated questions. First, what type of background knowledge, or prior, do participants have when they arrive in the experiment (i.e., in the very first task)? They could arrive with a naïve prior about the sequence of stimuli, or they could come with some structured prior background knowledge. Second, are participant’s experiences through the tasks changing how they approach subsequent tasks? To answer these questions, we first examined the distribution of choices over locations (aggregated over participants) for each of the first four predictions and each of the tasks, and we then quantitatively assessed the observed patterns.

To aid in the interpretation of the distribution of early behaviour from participants, we contrasted human behaviour against two reference artificial learning agents: (i) a naïve artificial learner and (ii) an ideal artificial learner, which are endowed with contrasting extremes of background knowledge in the task (see Methods). The naïve artificial learner has no knowledge of the generative family and instead uses count statistics to infer the transition probabilities. Thus, the first four predictions for the naïve artificial learner, which are made before the wolf revisits a location, are all random (i.e., because the agent starts with a flat prior with equal probability of choosing all locations, such an agent can only begin to make informed/non-random choices after a location has been visited at least once). At the other extreme of the spectrum is the ideal artificial learner. This artificial learner has perfect knowledge of the generative family and uses the observations to make inferences about which specific generative model controls the current stimulus sequence. While the first prediction must necessarily be a guess for this model, predictions become increasingly informed by the stimulus sequence from the second prediction onwards.

As for the first question: did participants come with a structured prior? Perhaps surprisingly, on the very first task, participants showed choices that were not only non-random, but were, if anything, somewhat aligned with the distribution of choices of the ideal artificial learner (task 1 in **Fig.2B**-**C**). This suggests that participants had some rough intuitions for what type of stimulus sequences were possible in the paradigm, or that the generative family we chose happened to broadly align with some default prior that the participants brought to the experiment. Regarding the second question: was there evidence of across-task learning? Inspecting the distribution of the first four predictions of the subsequent tasks revealed that participants’ background knowledge became increasingly more similar to the one from the ideal artificial learner (tasks 2, 3 and 4 in **Fig.2B-C**). This suggests that during their experience of a task, participants were able to change the background knowledge with which they would approach future tasks. In short, participants’ improvement of their background knowledge revealed across-task learning.

To quantitatively analyse the observed patterns, we derived a measure of early behaviour. As noted above, for each of the tasks, we started with the first four choices from participants and the expected distribution of choices from the two extreme agents in each task (see Methods). We then created an axis of background knowledge by using the expectation of choice probabilities from the two learning agents (the naïve and the ideal artificial learners). Projecting participants’ choices onto this axis has a natural interpretation: the highest values on the axis would be attained by choices that have the highest likelihood under the distribution of the ideal artificial learner and lowest likelihood under the distribution of the naïve artificial learner. However, while we have access to the choice probabilities of the artificial learning agents, we only have access to the concrete predictions from each participant. To allow for direct comparison of choices from humans and the artificial learning agents, we constructed a distribution of behaviour from choices of the artificial learning agents by sampling the first four predictions from the distribution of probability over choices for each of the two agents. Thus, for each human participant, we sampled a set of four predictions under each of the artificial learning agents’ distributions, we then projected the background knowledge from humans’ and agents’ choices onto the same axis, which allowed us to visualize comparable distributions of behaviour from participants and from the artificial learning agents (**Fig.2C**, left).

We first confirmed that participants start the experiment with priors that are not naïve but roughly aligned with the ideal artificial learner. Indeed, at the start of the first task, the distribution from humans (grey) was far from the distribution of the naïve learner (pink) and instead was shifted in the direction of the ideal artificial learner (blue). We used the area under the *receiver operator characteristic curve* (*AUC*, i.e., a non-parametric statistic), along with a permutation test in order to directly compare the statistical difference between distributions. The distribution of background knowledge from participants in the first task was far from that of the naïve artificial learners (*AUC* = .77, *p* < .001, see **Table 2**), and was also significantly lower than the distribution of the ideal artificial learners (although it was shifted in the direction of the ideal artificial learners; *AUC* = .04, *p* < .001). We then confirmed that participants improved their background knowledge through experience in the tasks, becoming increasingly similar to the distribution of the ideal artificial learners (**Fig.2C**, right). For this, we compared the background knowledge from participants across all contiguous pairs of tasks (i.e., task 1 *vs* task 2, task 2 *vs* task 3, and task 3 *vs* task 4). Crucially, the distribution of background knowledge was significantly higher (toward the ideal artificial learner) for task 2 compared to task 1 (*AUC* = .75, *p* < .001) and task 3 compared to task 2 (*AUC* = .56, *p* < .001), but did not differ statistically between task 3 and task 4 (*AUC* = .509, *p* > .4).

Importantly, our results are robust across a diverse set of experimental manipulations. For each and every one of the ten separate experiments that we ran, participants (i) arrived in the experiment with a structured prior, and (ii) showed an increase in background knowledge as they experienced the tasks of the generative family (see **Table 2**). This was the case not only when the tasks were delivered in the visual modality, where simple spatial priors may have existed (e.g., left to right, top to bottom visual scanning), but also when the tasks utilized an auditory stimulus, where spatial priors are arguably less likely (i.e., in 3 of our experiments, participants had to predict, instead of the location of the wolf, which of four syllables would be heard next). Together, our results show a strong tendency for participants to approach new learning problems in a structured way, making use of background knowledge acquired prior to the current task, and refining their background knowledge for their future learning experiences.

### Mixture-of-Generative-Models (MGM) learning explains within and across task behaviour

Yet, while the results above offer compelling evidence that human participants are learning the structure of the generative family across tasks, this result may seem difficult to reconcile with the overall choice performance shown in Fig.2A. How can participants be moving closer and closer to the optimal prior across tasks, yet be so far away from making optimal choices within given tasks (e.g., task 4 where there is the highest stochasticity)? We propose that the apparent discrepancy between (i) the highly structured early approach to a task, and (ii) the decreased performance observed in tasks with higher stochasticity, can be accounted for by assuming participants approach our tasks with a strong model-based approach. Specifically, we propose that participants start each novel task using a mixture-of-generative-models learning approach. In this framework, a pool of candidate generative models (which increasingly resembles the broad generative family as participants move through the tasks) is arbitrated against each other with regard to which best explains the stimulus sequence. Below we describe in depth (i) details of this framework, (ii) how it accounts for the results, and (iii) the novel predictions that it makes.

Multitask learning is a computationally hard problem. In our study, there is potentially an infinite number of generative models that could have generated each of the task sequences. A sensible strategy in this context is to use a mixture-of-generative-models (MGM) learning strategy, which combines elements from mixtures of models for time series (Fox et al., 2009; Fox & Jordan, 2013) as well as the intuitive biases of probabilistic programming (Tenenbaum et al., 2006; Tenenbaum & Griffiths, 2003). Under MGM learning, participants simultaneously entertain a number of candidate generative models at the start of each task. Then, as the stimulus sequence unfolds, they arbitrate the models against each other. Generative models that make correct predictions gain credit (i.e., participants increase their confidence in that model being the one controlling the stimulus) and thus are more likely to guide a participant’s future choices, while generative models that do not make correct predictions tend to lose credit and fade into the background (i.e., no longer influence choices). MGM learning is ideally suited to model learning in our study given the paucity of information participants were given to complete the task (see supplementary materials for a precise description of the prediction and production task instructions). Indeed, at no point were the participants instructed that a single generative model operated in each task. Here it is important to note that although in the case of the experiment at hand a single generative model operated in each task, if participants had assumed that a single model controlled the stimulus and this was not true (e.g., if a task had been controlled by a mixture of several generative models) it could lead to significant learning failures. Specifically, participants’ internal model in such a scenario, would tend, after a few switches in the generative models, to predict all transitions with equal likelihood. Conversely, an MGM learning strategy would enable the correct inferences to be made as time unfolds. Our use of an MGM learning framework is further motivated by the fact that the early task behaviour observed above is naturally incorporated in such a framework, in addition to a host of other in-task choice behaviours that we examine below.

Participants more likely guide their choices using the generative model for which they have higher confidence. Specifically, under the MGM framework, participants’ confidence in each of the models, that is, the probability with which they think each model is controlling the stimulus, is constantly updated with new stimulus observations (**Fig.3A**). Participants need to simultaneously (i) arbitrate amongst the generative models they entertain in mind, and (ii) learn their parameters (e.g., estimate the dominant transition probability values for estimating the likelihood of an observation under each of their candidate models). Amongst the candidate generative models that participants entertain, one model predicts with a high probability the most likely stimulus on every trial, in agreement with the true generative model (albeit perhaps with a different dominant transition probability value). Henceforth, we call this the “dominant model”, which is characterized by having the dominant pattern that matches the dominant pattern of the true generative model (note that it is referred to the dominant model regardless of whether participants are actually using it to guide their choices at any given time). The dominant model thus predicts with the highest probability each of the dominant transitions, that is, the most likely next locations given any location, but predicts with low probability each of the non-dominant transitions. In other words, because of the stochasticity in the stimulus sequences, as sampled from the true generative model, the stimulus sometimes violates the dominant pattern (i.e., the stimulus does not always transition to the most likely next location). Alternative “non-dominant models” predict the non-dominant transitions with a higher probability than the dominant model, and participants may use those alternative models to guide some of their predictions, which will then result in non-dominant choices. Thus, under the framework, there is an important interplay between (i) evidence provided by the stimulus sequence which can be either in favour or against the dominant model, and (ii) the credit that participants assigned to the dominant model and which can be inferred by the type of choices they make (i.e., choosing the most likely next location or not), and which will determine how much participants may update each model’s parameters.

**Fig 3.**
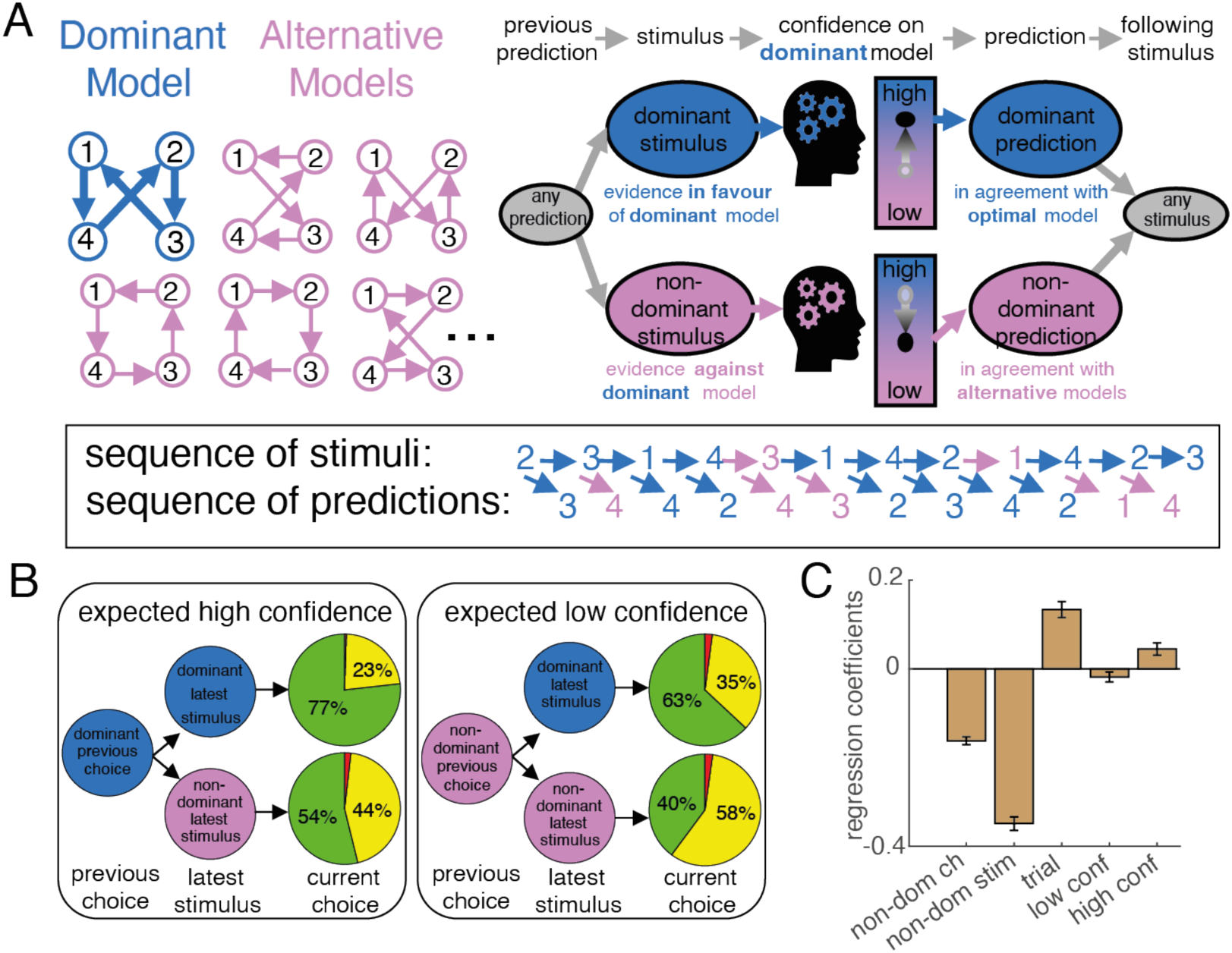
Learning via a mixture-of-generative-models. (**A**) ***Arbitrating amongst a pool of generative models***: Under the MGM framework, participants entertain a pool of candidate generative models: the dominant model (i.e., the one with the correct dominant pattern, blue) but also other alternative models (i.e., models with different dominant patterns, pink). As the stimulus sequence unfolds, stimulus transitions either give evidence in favour of the dominant model (i.e., when the participant observes a dominant stimulus transition, blue) or against it and in favour of some of the alternative models (i.e., when the participant observes a non-dominant stimulus transition, pink). The observation of new stimulus transitions thus causes fluctuations in participants’ belief about which model controls the stimulus sequence. As confidence in the dominant model increases, so does the probability of participants using that model to guide their choices—and thus the probability with which they will make a dominant prediction (i.e., predicting the dominant transition; blue). When evidence for the dominant model decreases (low confidence), alternative models are more likely to guide participants’ choices and thus result in a non-dominant prediction (pink). In short, the “errors” that participants make (i.e., when they make a choice to a location other than the one that would be suggested by the dominant transition) should not be randomly distributed in the sequence. Instead, the probability of certain “errors” should depend upon whether a non-dominant transition was observed (or a non-dominant choice was made on a previous trial). (**B**) ***Participant errors are structured according to MGM:*** Participants’ choice behaviour indicated that the proportion of dominant choices in one trial (green) depended on the recent history leading up to that trial. As predicted by MGM learning: when the latest stimulus was dominant, participants were more likely to use the dominant model—and thus make a dominant prediction—as compared to when that stimulus was non-dominant. Similarly, when the previous choice was dominant, the next choice was more likely to be dominant as compared to when the previous choice was non-dominant (i.e., a choice revealing that participants were not relying on the dominant model at the time). (**C**) ***Evidence for gated learning:*** We ran a logistic regression analysis, whereby we predicted the trial-by-trial choice type of each participant (i.e., dominant or non-dominant, 1 or 0) as a function of five predictors. The first predictor, whether the previous choice was non-dominant (*non-dom ch*; 1 or −1), helped us confirm that performance was lower after having made a previous non-dominant choice. The second predictor, whether the latest stimulus was non-dominant (*non-dom stim*; 1 or −1), helped us confirm that performance was lower after having observed a non-dominant stimulus. The third predictor, an index of time elapsed (*trial*; z-score of trial number) helped us capture the general tendency for performance to improve with time. The fourth and fifth predictors are interaction terms between the latest stimulus type (non-dominant or not, 1 and −1) and time elapsed (z-score of trial index) but separately for trials when participants have a predicted low confidence on the dominant model being in control of the stimulus (fourth predictor, *low conf*) and when they have a predicted high confidence (fifth predictor, *high conf*).These last two predictors provide evidence for gated learning, which predicts that participants learn to deal with (i.e., learn to have their performance unaffected by) non-dominant stimuli mostly when they experienced strong evidence (high confidence) of the stimulus being controlled by the dominant model. The reported coefficients have arbitrary units (but all predictors have close to unit standard error), we report the mean and SEM across participants.

The MGM framework can explain both within- and across-task patterns of behaviour. The across-task learning results, interpreted within the MGM framework, suggest that the pool of candidate generative models that participants use is increasingly more similar to the pool of generative models within the generative family. In line with this, only 1.3% (± .07% SEM across participants) of choices in the fourth task fell outside the generative family; the percentage of choices was computed for each participant using all choices and we report the mean and standard error of the mean across participants. Similarly, the drop in within-task learning performance that we observed for tasks with higher stimulus stochasticity has a natural interpretation under MGM learning. Specifically, more stochastic stimulus sequences are compatible with a larger number of generative models, making the arbitration harder and explaining why participants may use alternative models to guide a higher proportion of their choices. Interestingly, typical explanations for the decreased performance for more stochastic sequences require invoking various suboptimal mechanisms (e.g., probability matching), which often implicitly assume that errors are randomly distributed across trials. Instead, MGM learning makes specific predictions about the structure of errors. Namely, error rates are expected to increase when (i) the most recent stimulus transition was non-dominant, as this casts doubt on the dominant generative model, and (ii) when the previous choice was non-dominant, because it reveals participants were not using the dominant model to guide their predictions, i.e., an alternative model had higher credit at the time.

To demonstrate that participants choices were well matched to this framework, we analysed the proportion of dominant choices made in groups of trials segregated according to the type of previous choice and latest stimulus type (**Fig.3B**). As predicted under MGM, a non-dominant stimulus had the effect of reducing the proportion of dominant next choices by ≈20% ± 1.3% SEM compared to a dominant stimulus (from ≈73% down to ≈53%). This reduction was observed regardless of the previous choice type (after a dominant choice, ≈22.9% ± 1.3% SEM; from ≈77% down to ≈54%; and after a non-dominant choice, ≈23.1% ± 1.3% SEM; from ≈63% down to ≈40%). Similarly, a previous non-dominant choice was followed by ≈10% ± 1.3% SEM fewer dominant choices compared to a previous dominant choice (from ≈70% down to ≈59%) and this was true regardless of the stimulus type (after a dominant stimulus, 13.7% ± 1.2% SEM; from ≈77% down to ≈63%) and after a non-dominant stimulus, ≈13.9% ± 1.4% SEM; from ≈54% down to ≈40%). This latter result, namely, the negative effect of a previous non-dominant choice, can be interpreted under MGM as reflecting participants’ decreased confidence in the dominant model, therefore increasing the probability of using an alternative model for guiding the following choice. Together, these results are highly consistent with the MGM view that participants’ confidence in their candidate generative models (for the most part within the generative family), fluctuates along with the evidence, as the stimulus sequence unfolds.

Another prediction from MGM is that learning the parameters of each model should be gated by confidence in the model. Confidence in the dominant model decreases when the stimulus violates the expectations from the dominant model, which in turn often leads to a non-dominant choice. As such, a low level of confidence in the dominant model can be inferred by observing a non-dominant choice. However, the influence of confidence on the dominant model is not limited to explaining the structure of errors. In fact, a crucial prediction of the mixture-of-models framework is that feedback, that is the observation of a stimulus in the sequence, needs to be attributed to the different models in proportion to how much each model is deemed responsible for generating the observation. A direct prediction of this process is that *learning should be gated by confidence*. Specifically, we predict that learning the **α** parameter (i.e., the dominant transition probability value) of the dominant model should be facilitated when confidence on the model is high compared to when it is low. In other words, participants need to learn to expect non-dominant transitions within the dominant model, but they are less likely to learn about them when they believe the sequence is being controlled by a different model.

We thus tested for the existence of this gated learning mechanism. For this, we ran a logistic regression with five competitive predictors (where the order of the predictors does not change the estimates of the regression coefficients). First, three predictors of the regression allowed us to control for the effects of the previous choice type, and latest stimulus type, as well as a rough estimate of the general effect of time on performance, then two predictors represented the two contrasting states of learning, that is, learning when confidence is high, and learning when confidence is low (**Fig.3C**). Indeed, and as expected from the MGM framework, and confirming results from the patterns of errors, we found a negative impact on performance (i.e., an increased probability of making a non-dominant choice) of having made a non-dominant choice in the previous trial (*t*(598) = −18.5, *p* < .001) and of observing a non-dominant transition with the latest stimulus (*t*(598) = −23.0, *p* < .001). Furthermore, this analysis showed a positive effect of time on performance (*t*(598) = 7.4, *p* < .001), i.e., on average performance improves over time.

Evidence for gated learning came from the remaining two predictors of the regression, which together probe in what conditions participants learned to deal with non-dominant stimuli. Specifically, non-dominant stimuli provide evidence that favours alternative models and have the effect of increasing the probability of non-dominant choices (i.e., participants are more likely to use an alternative model to guide their choices). However, under MGM, participants can gradually learn the correct dominant transition probability value, and hence, also the probability values of the non-dominant transitions; with more accurate estimates of the non-dominant transitions, participants will, on average, be less surprised by non-dominant stimuli (i.e., with accurate estimates, it is less likely that non-dominant transitions appear to be too frequent or too rare given the generative model). In short, under MGM we expect participants to learn to be less surprised and less affected by non-dominant transitions. Consistent with this, we recovered a positive effect of “learning to deal with non-dominant stimulus transitions” on performance for cases when participants were expected to have high confidence in the dominant model (fifth predictor, *high conf*, *t*(598) = 3.19, *p* < .01). When participants were predicted to have been using an alternative model, the effect was lower than for high confidence (fifth vs fourth predictors, *high conf* vs *low conf*, *t*(598) = −5.33, *p* < .001) and not different from zero (fourth predictor, *low conf*, *t*(598) = −1.62, *p* = .1), showing that learning was gated by confidence. These gated learning effects would be otherwise hard to explain, but are easily and naturally explained under MGM learning, providing a final confirmation that indeed participants are engaging in a mixture-of-generative models learning strategy to solve our multi-task experiment.

Finally, while the structured pattern of errors provides strong evidence in favour of the MGM framework, it remains possible that simpler forms of learning, like trial-and-error learning could contribute in parallel to control behaviour. Such simpler learning models predict that rewarded choices increase the probability of repeating the choice in the future. We thus sought to assess the impact of reward on subsequent behaviour. However, in our experiment the effect of reward (i.e., catching or not catching the wolf) is confounded with the effect of the previous prediction type (dominant or not) and the latest stimulus transition (dominant or not) across all trials. To avoid the confound, we focused only on those trials where participants made non-dominant choices and the stimulus that followed was also non-dominant. Because in this circumstance the choice could either be rewarded or not (i.e., in 50% of trials where the participant made a non-dominant choice *and* the wolf moved to a non-dominant location, the wolf would actually be caught and in the other half of trials the wolf would not be caught) this confound was eliminated. Learning by trial-and-error would predict that rewarding non-dominant choices (i.e., when the wolf was caught) should increase the probability of future non-dominant choices (and thus decrease the probability of dominant choices) compared to not rewarding non-dominant choices (i.e., when the wolf was not caught). Instead, we found that reward on its own had only a minor impact on participants’ predictions. The percentage of dominant predictions was 42.3% (± 1.1% SEM) after a rewarded non-dominant choice and 39.8% (± 1.1% SEM) after an unrewarded non-dominant choice; this small change (2.5% ± 1.4 SEM) is, if anything, going in the opposite direction of what would be expected from trial-and-error learning. This analysis suggests that the effect of simple trial-and-error learning, in our experiment, is negligible.

## Discussion

How humans and other animals learn a generative model for a single task has been a topic receiving ample attention. Yet, there is limited knowledge about the role that generative models themselves play in guiding future behaviour. Here, we presented a novel experimental paradigm where human participants went through a series of prediction tasks (i.e., on each trial they explicitly predicted where a stimulus would appear next). Critically, the generative models that controlled the stimulus sequences in each of the prediction tasks were sampled from a common generative family: a space of possible generative models respecting a set of rules. This multi-task prediction learning paradigm allowed us to investigate not only within-task learning but also across-task learning, and thus address a number of key questions about the role that generative models play during learning.

Humans have been shown to approach learning problems in perceptual, decision-making and cognitive everyday tasks, equipped with inductive biases. These biases come in the form of prior knowledge and intuitions, which can be both innate and learned, that exploit structural properties shared across different tasks, and which result in learning in a particular task to unfold more quickly than would otherwise be expected (Acuña & Schrater, 2010; Braun et al., 2010; Chomsky, 1980; Harlow, 1949; Lake et al., 2017; Tenenbaum et al., 2011). The combination of our multi-task prediction learning paradigm with the analysis of the very early behaviour allowed us to determine that participants came to the experiment with good intuitions (in the form of structured priors) which they rapidly updated to better match the true generative model of the tasks. Our findings thus suggest that structured priors are a key ingredient for bringing about inductive biases.

Our results extend our current knowledge of inductive biases and structure learning in important ways. Structure learning is often conceptualised as a reduction of the dimensionality of the space over which to search for a solution to the new task that causes learning to occur faster (Braun et al., 2010) and many previous reports suggest that structure learning arises slowly over repeated exposure to (often several) different instances of tasks within the task space. For example, Harlow’s research on the formation of learning sets required tens or sometimes hundreds of learning problems (Harlow, 1949). In contrast, here we observed that exposure to a single task allowed our participants to improve their knowledge of the task space. Second, while previous studies either focused on learning of multiple simple tasks (Kattner et al., 2017; Schulz et al., 2020), or learning of a single complex graph-like structure (Cleeremans & McClelland, 1991; Garvert et al., 2017; Schapiro et al., 2013), we focused on how humans learn multiple complex structures, an issue which had not been looked at until very recently (Mark et al., 2020; Wu et al., 2019). Third, studies typically focus on the consequences of what is being transferred, for instance, whether there is an immediate benefit to performance or a change in the rate of learning (Braun et al., 2010; Kattner et al., 2017); our paradigm gives us additional insights into the *content* that is being transferred, specifically, the pool of candidate models that are used for explaining the task increasingly matches the true pool of possible models within the paradigm. Fourth, previous studies of generalization have primarily focused on how learners search for solutions (i.e., a parameter set that is incrementally tuned) that help them best explain the new task. Instead, we propose that participants faced with complex multi-task environments likely engage in a mixture-of-generative-models (MGM) learning strategy, whereby they maintain a multitude of competing solutions which are continuously updated and mixed in order to form predictions.

Behaviour under an MGM learning framework has all the key characteristics of our human learners. Humans seem to simultaneously engage not only in (i) structure learning, i.e., the discovery of the possible structures at play, meaning both discovering them from scratch as well as discovering that some previously learnt structures are useful in the current context, and in (ii) parameter learning, i.e., the fine tuning of the parameters of any one structure, but also in (iii) the arbitration amongst different competing structures in any one task, a process which could be called structure switching. Finally, previous studies that have attempted to capture the variability of choices in learning tasks through the lens of optimality, have done so by casting the problem in terms of balancing the trade-off between (i) the benefits of exploiting the best candidate structure thus far and (ii) the costs of exploring the possible other structures (Acuña & Schrater, 2010; Gershman, 2018). However, in our tasks, such a trade-off is non-existent because the observations are independent of the choices. Instead, we propose that MGM learning is able to explain variability in choices as a natural consequence of the variability in the priors that participants bring to the experiment. Additional evidence in support for the MGM framework comes from the patterns of gated learning observed from participants’ behaviour which are highly diagnostic of participants’ usage of the strategy.

The MGM framework is a fully Bayesian forecast motivated by work in machine learning and artificial intelligence. It is a system with strong inductive biases that nonetheless allow cross-task generalization, and as such, it is more consonant with a probabilistic programming approach (Tenenbaum et al., 2006; Tenenbaum & Griffiths, 2003) than deep learning approaches which do not require inductive biases but instead have much higher requirements of data (Wang et al., 2018). A strength of the MGM framework is that it helps solve two problems simultaneously: first, how to allow strong inductive biases while preserving enough flexibility to deal with the idiosyncrasies of the environment, second, how to mimic the fast learning that humans are capable of, while also capturing the individual differences in learning speeds. The framework thus relies on hierarchical generative model families which exploit compositionality (i.e., models factor into reusable parts) — allowing flexible mapping between an abstract dynamics space and observation spaces (i.e., factoring the dynamics and the observation). The framework also relies on the minimum entropy principle: learners have a predisposition to hasten their learning, by having a bias for determinism (Brand, 1999), at the risk of learning collapse in stochastic domains, where a maximum entropy assumption would not lead to such collapse (Jaynes, 1982). Preserving flexibility while using strong inductive biases is a serious theoretical and computational challenge, which humans seem extremely good at achieving. We propose that humans rely on MGM as a strategy to solve this challenge.

Finally, the MGM framework not only provides a single, theoretically sound, explanation that accounts for all those features of the data without the need to invoke any suboptimal mechanisms on the side of participants, but it also makes important predictions about the behavioural consequences of manipulating participants’ priors that could guide future lines of research. Two specific aspects of participants’ prior are predicted to be of particular importance: (i) the size of the model pool (i.e., the number of candidate models that participants are arbitrating against one another), and (ii) the bias for determinism (i.e., the amount of stochasticity that participants allow their candidate models to have at the start). Regarding the size of the pool of models, there are pros and cons. If participants arrive at the task with a large number of competing models, they are more likely to have a model that is good for solving the task. However, a larger pool of models also means that the space of inferences, and the time taken to find a good model, grows exponentially with the number of candidate models. Regarding the bias for determinism, the more deterministic a model is, the less responsibility it has for explaining transitions that deviate from its predictions, and thus, the less it will be updated to incorporate more stochasticity (as long as there are other models that explain those transitions). This latter point can have disastrous consequences for performance: having too strong a prior for determinism can lead participants to never learn to expect non-dominant transitions within the dominant model, and instead be stuck in a loop where they constantly switch between deterministic models that capture regularities at a short temporal scale. Interestingly, however, the inductive biases we see, including the bias for determinism, reveal a strong desire from our participants to learn quickly. Getting a better understanding of how to influence participants’ inductive biases could help capitalize on participants’ desire for learning when facing new learning problems. These insights could benefit fields as diverse as artificial intelligence and educational sciences.

## Acknowledgements

We want to thank Aaron Cochrane and Ornela De Gasperin for useful comments on the manuscript, and Amanda Yung for help with programming the tasks. This work was supported and funded by: a Swiss National Science Foundation grant: 100014_15906/1 to DB; the Luxembourg National Research Fund: ATTRACT/2016/ID/11242114/DIGILEARN to PCL; the INTER Mobility/2017-2/ID/11765868/ULALA to PCL and PS. The Office of Naval Research grant N00014-17-1-2049 to CSG. The European Union’s Horizon 2020 research and innovation program under Marie Skłodowska-Curie grant no. 661667, Learning Determinants to IA. The Office of Naval Research grant MURI GRANT N00014-07-1-0937 to DB.

## Author contributions

PCL, CSG, PS and DB conceptualized the research idea and created the research design; SHC and PCL programmed the experiments. SHC, PC, and IA collected the data; SHC, PCL, CSG, PS and DB designed the analyses; SHC, PCL, and PS analysed the data; SHC and PS developed the models; SHC, PCL, CSG, PS and DB interpreted the results; SHC drafted the manuscript; SHC, PCL, CSG, PS and DB finalized the manuscript; SHC and PCL managed the submission process. All authors approved the final manuscript.

## Methods

### Ethics and consent

The study was approved by the Ethical Review Panel of the Faculty of Psychology and Educational Sciences of the University of Geneva. All participants were recruited using Amazon’s Mechanical Turk (MTurk) platform. Participants were all based in the US, at least 18 years old and had an approval rating of at least 95%. After reading information about the study, participants were asked to consent by clicking on a checkbox.

### The Catch The Wolf paradigm

Catch The Wolf is a multi-task learning paradigm. It presents participants with a series of four different sequential *prediction tasks*. In each prediction task, participants observed a wolf successively appear in one of four possible screen locations and they were tasked to continuously predict where the wolf will appear next. After each prediction task, participants performed a brief *production task*, where they were asked to mimic the behaviour of the wolf they just experienced in the latest prediction task by producing a sequence of locations. Crucially, the behaviour of the wolf within a particular task is generated by a specific generative model (i.e., a specific location-to-location transition matrix). While each of the four tasks can be determined by a different generative model—and hence requires participants to learn each task anew—all generative models stem from a common generative family—knowledge about this generative family could be exploited when confronted with new generative models.

#### The generative model

The sequence of wolf locations in a given prediction task is generated as a first Order Markov chain, where the location of the wolf in one trial depends solely and probabilistically on the location from the previous trial. This generative model can be described by a transition probability matrix from one trial’s location to the next trial’s location. However, not all transition probability matrices are valid under the generative family: valid transition matrices need to respect five rules. First, there is one maximum value per row which we call the “dominant transition”, i.e. from any one location there is a unique location which is the most likely next location. Second, all diagonal values are set to zero, i.e. there are no self-transitions, or the wolf never reappears in the same location from where it just disappeared. Third, there is a uniform stationary distribution, i.e. on average, all locations are visited by the wolf with the same frequency. Fourth, the value of the dominant transition probabilities is equal across four rows, i.e. there is a single dominant transition probability value to be learnt. Fifth, all non-dominant and non-diagonal transitions have the same probability value, i.e. there is a single non-dominant transition probability value.

These five rules strongly constrain the space of possible wolf behaviours. In essence, given that we have four locations for the wolf to appear, this space of valid transition matrices can be characterized by two variables. First, the dominant transitions of valid matrices follow one of six possible patterns (i.e., these patterns indicate which is the most likely next location from any one location) of which we used only four in our studies. Second, valid transition matrices are characterized by the value of the dominant transition probability (i.e., the probability with which the wolf will move to the most likely next position). Thus, the transition probability matrix controlling the wolf on any one task was obtained by selecting (i) a dominant transition pattern and a (ii) dominant probability value (**Fig.1A**).

#### Prediction tasks

Across most experiments the basic paradigm consisted of four prediction tasks that participants completed in succession (**Fig.1A**). In each of the four prediction tasks, participants observed a wolf (a cartoon picture of the head of a wolf) that successively appeared in either one of four possible screen locations (**Fig.1B**). Participants’ goal was to continuously predict (i.e., one prediction for each trial) the wolf’s next location. The wolf’s behaviour can be fully described by a transition probability matrix that governs the probability that the wolf will move from any one location to any other location. To perform well, participants had to learn (and select) the most likely destination for each possible current location of the wolf. The sequence of transitions was strictly identical for each participant within an experiment. For each Prediction Task these sequences were generated in such a way that the empirical transition frequencies approximately matched the transition matrix of the true model.

The true model (described in detail in the following section) can be described as consisting of two parts: (i) a permutation matrix (which unique location is the most likely from each position) and (ii) a dominant transition probability (i.e., the probability with which the wolf will move onto the most likely location). Because participants find it easier to learn more deterministic sequences and to ease participants into the experiment, we implemented a scaffolding strategy whereby the value of the dominant transition probability was gradually decreased across tasks (but remained constant within a task). Thus, for most experiments the dominant transition probability (**α**) for prediction task 1 had a value of 1.0, for prediction task 2 it had a value of 0.95, for prediction task 3 it had a value of 0.85 and for prediction task 4 it had a value of 0.75. Similarly, the number of trials was set for each task in such a way as to keep the task duration as short as possible while allowing enough trials for most participants to achieve learning.

#### Production tasks

Immediately after each Prediction Task, participants were asked to imitate the wolf’s stochastic behaviour by sequentially clicking on one of the four possible locations. The exact instructions stated the following: “You’ve observed how this wolf behaves. Your goal now is to be the wolf. Click on the target locations to make the wolf appear just like the one you’ve just observed.” They were asked to make 24 clicks creating 23 wolf moves. The production tasks not only helped emphasize the contextual cues that one prediction task was over, but they also provided us with a different perspective to investigate what participants had learnt about the behaviour of the wolf during the prediction task.

### Experimental manipulations

We collected data across 854 participants over a total of ten experiments. All participants within an experiment experienced the exact same sequence of stimuli. However, we manipulated the stimulus sequence across experiments in important ways (see **Table 1**). First, in separate experiments, stimuli were presented in the visual and auditory modalities respectively. In the “Catch The Wolf” experiments the stimulus was visual (the cartoon face of a wolf; as described above) and could appear in one of four possible locations. Participants made their predictions (and productions) by clicking on either one of the four locations. In the “Catch The Sound” experiments the stimulus was auditory (brief audio-tracks of the syllables “bi”,”da”,”fu” and “go”) and participants made their predictions (and productions) by clicking on circular placeholders with the corresponding text (i.e. “bi”,”da”,”fu” and “go”) whose relative spatial order on the screen varied randomly from trial to trial. Second, different experiments had a different progression of tasks. With each task being described by: (i) number of trials, (ii) the identity of the dominant pattern and (iii) the dominant transition probability value; (see **Fig.1A**).

**Table 1.** Experiment descriptions.

| Exp ID | Number of Participant | Dominant patterns (Pat), dominant transition probability value (in |  |  |  | Sensory modality |
| --- | --- | --- | --- | --- | --- | --- |
|  |  | Task 1 | Task 2 | Task 3 | Task 4 |  |
| 1 | 173 | Pat1 100% (76) | Pat2 95% (101) | Pat3 85% (151) | Pat4 75% (201) | visual |
| 2 | 222 | Pat1 100% (50) | Pat2 95% (75) | Pat3 85% (100) | Pat4 75% (200) | auditory |
| 3 | 64 | Pat1 100% (76) | Pat1 85% (101) | Pat1 85% (151) | Pat4 75% (201) | visual |
| 4 | 71 | Pat1 100% (76) | Pat2 85% (101) | Pat3 85% (151) | Pat4 75% (201) | visual |
| 5 | 48 | Pat3 100% (76) | Pat3 95% (101) | Pat3 85% (151) | Pat4 75% (201) | visual |
| 6 | 44 | Pat1 100% (76) | Pat2 95% (101) | Pat3 85% (151) | Pat4 75% (201) | visual |
| 7 | 63 | Pat3 100% (76) | Pat3 95% (101) | Pat3 85% (151) | Pat4 75% (201) | auditory |
| 8 | 56 | Pat1 100% (76) | Pat2 95% (101) | Pat3 85% (151) | Pat4 75% (201) | auditory |
| 9 | 56 | Pat1 100% (76) | Pat3 95% (101) | Pat4 75% (201) | only 3 tasks | visual |
| 10 | 57 | Pat1 100% (76) | Pat4 75% (201) | Pat3 85% (151) | only 3 tasks | visual |

### Exclusion criteria

Recruiting participants on online platforms improves the convenience of collecting data from a larger pool of participants. However, it means we have to pay attention to only include data from participants who are: (i) completing the experiment, (ii) following the instructions adequately and (iii) trying hard to do the tasks. We thus decided to impose the following exclusion criteria to filter out participants that were not actually doing the task. First, we decided to exclude 393 participants who failed to provide a response in at least 85% of the trials of any one prediction task (most of these are entries from participants who started the experiment but soon after abandoned it). Second, we decided to further exclude the 26 participants whose performance (i.e., proportion of choices in agreement with the true generative model) was at or below 25% (i.e., chance level) at any one prediction task. Third, we excluded a further 124 participants who produced 5 or more consecutive self-transitions (i.e., simply clicking the same position over again) in any one production task. In total, 854 participants survived all exclusion criteria.

### Performance progression in relation to the true generative model

In order to assess within-task learning, we decided to compute performance in relation to the true generative model: the proportion of choices that match what an agent with full knowledge of the true generative model (an agent that does not need to learn). We then aggregate the proportion of those choices for each participant made within time bins of 25 trials in length. The learning curves shown in **Fig.2A** are obtained by showing the mean and standard error of the mean of choice performance across participants for each time bin and for each task.

### Artificial learning agents

As a tool for analysing the quality of participants’ background knowledge and how it changed as participants experienced the different tasks, we created an axis that spanned the space of possible knowledge quality about the generative knowledge, going from zero knowledge of the generative model all the way to perfect knowledge of the generative model. The two extremes of this axis were obtained contrasting the choice probabilities of two artificial learning agents that we describe next: (i) a naïve artificial learner and (ii) an ideal artificial learner. First, we describe a naïve artificial learner who comes with a flat prior about what could happen in each of the tasks and simply keeps track of the observed transitions so that the next time a location is visited, it chooses the transition with the highest number of observations (with a random choice amongst any locations that have tied counts). Second, we describe an ideal artificial learner who knows the generative family and makes inferences about which specific generative model governs the current stimulus sequence and makes optimal inferences about the most likely next transition. Specifically, the ideal artificial learner starts with a flat prior over the possible transition matrices within the high-level generative model and updates its belief upon every new observation to make inferences about (i) the probability of that each of the six possible dominant patterns is governing the stimulus, (ii) the value of the dominant transition probability. By marginalizing over these two dimensions, the agent can then select the most likely next location given the observed sequence thus far.

### Quantifying high-level generative model knowledge (early behaviour analysis)

Priors can quickly change in the face of new evidence. To analyse the type of background knowledge that people came with at the start of any one task, we decided to use the behaviour of the first four trials of each prediction task. The first four trials are special because each of the predictions in those trials is done before the wolf has revisited a location, and thus choices are entirely driven by the prior with which participants arrive at the task and not by the evidence they have acquired in the current task. Note that this need not mean either that the first four locations visited follow the dominant pattern of the task, or that the wolf on the fourth trial did not reappear in a location it had already visited (because the fourth choice is done before the wolf appears for the fourth time). For example, in task 4 of experiments 1-8, the first four stimulus transitions are not following the dominant pattern: the first stimulus appears in the top right location, under the dominant pattern it should go to the bottom right, but instead went to the top left location on the second trial. This shows that those first four trials help dissociate the choices that would be correct under the true generative model, from the choices that an ideal learner would make given the evidence that had been observed to that point.

To make sense of participants’ early behaviour in the tasks we decided to contrast it against the expected behaviour of the two benchmark learning models described above. Specifically, for any one trial we derived the probability of observing the choice of any one participant given any one of the two models. This allows us to obtain a measure of the relative probability that a participant’s prediction in the i^th^ trial resembles the expected prediction under the ideal artificial learner or under the naïve artificial learner:

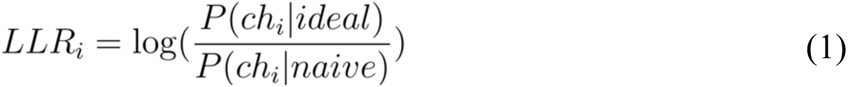

However, because not all trials are equally diagnostic of the difference between the two models, we first computed the diagnosticity of each trial by computing, for each of the trials, how divergent the probability distributions of predictions were for the two models. We thus computed the Kullback-Leibler divergence (KL divergence) between the two distributions. Because the KL divergence is an asymmetric measure, we computed the divergence using each of the distributions as a base and summed the divergences to obtain our diagnosticity measure for each trial:

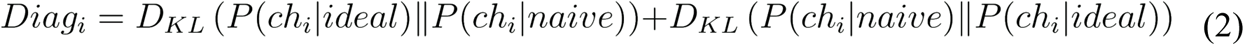

Finally, we obtain a single measure of each participant’s early behaviour in each task by computing the diagnosticity-weighted average log-likelihood ratio, *dwLLR*:

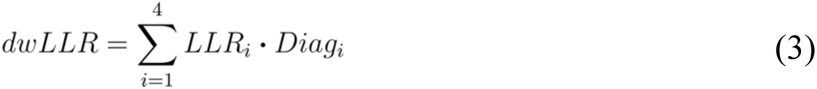

Using this measure, a distribution of behaviour for all participants in each of the prediction tasks can be retrieved. Importantly, the axis over which participants’ individual choices are projected is constructed using the theoretical expectation of choice probabilities under the two benchmark models. This way, the two extremes of the axis correspond to choices that are maximally discriminant of the two models (which need not be the most typical choices of either model). Instead of simply obtaining the distribution of the *dwLLR* measure computed from participants’ choices, we decided to compute a similar distribution from choices sampled from the two benchmark models. Thus, for each actual participant, we simulated one instantiation of each of the two benchmark models for each of the trials and constructed a distribution of *dwLLR* for each model and each task (see **Fig.2C, grey**). More specifically, for each of the first four trials, each of the benchmark models describes a probability distribution over choices. For each participant we decided to take one instantiation of choice (i.e., taking a random draw from the probability distribution) for each trial, under each of the two models. This allowed to create two separate pools of simulated agents (and their choices) for which we also computed the *dwLLR* measure (see **Fig.2C, pink and blue**).

In order to test whether the distribution of background knowledge from participants in task 1 differed statistically from the distribution of the naïve artificial learner, we computed the area under the curve (*AUC*) of a receiver operator characteristic score along with a permutations test (with 1000 reshuffles) in order to obtain a null distribution of the differences in score and p-value (i.e., the proportion of the 1000 reshuffles that had an *AUC* score as extreme or more extreme than the unshuffled score). We used the same statistical procedure to test whether the distribution of background knowledge differed between: (i) task 1 and task 2, (ii) task 2 and task 3, and (iii) task 3 and task 4. The results of these tests for each experiment as well as aggregating participants over all experiments, is concentrated in **Table 2**.

**Table 2.** Statistical comparison of the Background Knowledge Distributions.

|  | task 1 > naïve |  | task 2 > task 1 |  | task 3 > task 2 |  | task 4 > task 3 |  |
| --- | --- | --- | --- | --- | --- | --- | --- | --- |
| **Experiment** | *AUC* | *p-val* | *AUC* | *p-val* | *AUC* | *p-val* | *AUC* | *p-val* |
| 1 | **.832** | **.000** | **.754** | **.000** | **.587** | **.002** | .509 | .757 |
| 2 | **.752** | **.000** | **.687** | **.000** | **.574** | **.001** | .507 | .738 |
| 3 | **.838** | **.000** | **.711** | **.000** | .586 | .107 | .555 | .182 |
| 4 | **.869** | **.000** | **.711** | **.000** | **.606** | **.015** | .493 | .926 |
| 5 | **.872** | **.000** | **.844** | **.000** | .552 | .253 | .458 | .268 |
| 6 | **.775** | **.000** | **.727** | **.001** | .557 | .400 | .591 | .126 |
| 7 | .565 | .196 | **.810** | **.000** | **.635** | **.000** | .492 | .772 |
| 8 | .575 | .169 | **.911** | **.000** | .545 | .317 | .482 | .690 |
| 9 | **.784** | **.000** | **.759** | **.000** | .500 | .999 | .500 | N/A |
| 10 | **.817** | **.000** | **.781** | **.000** | .465 | .540 | .500 | N/A |
| 11 | **.762** | **.000** | **.800** | **.001** | .529 | .733 | .514 | .913 |
| All | **.771** | **.000** | **.752** | **.000** | **.567** | **.000** | .509 | .427 |

### Finding structure in participants’ errors

The MGM framework predicts that errors in the tasks would not be random but that they should be structured and predictable given the trial history. We thus decided to look at the proportion of each choice type: (i) dominant (i.e., the choice is optimal under the true generative model), non-dominant (i.e., the choice is not the optimal choice under the true generative model, but it could be guided by an alternative model within the generative family) and (iii) self-transition (i.e., predicting that the wolf will appear where it just disappeared, guided by a model outside the generative family).

The proportion of choices of each type was examined conditional on history or recent previous events, in line with predictions of the MGM framework. Specifically, both (i) the type of latest stimulus (i.e., whether the latest stimulus transition was dominant under the true generative model or whether it was non-dominant), and (ii) the type of previous choice (i.e., whether the previous choice was dominant under the true generative model or not), exerted an influence on choices.

The rationale behind grouping trials based on the recent history leading to each of them is that the MGM framework makes explicit predictions about how the proportion of current choices should change as a function of last stimulus and choice. For instance, when the stimulus is non-dominant under the true generative model, it casts doubt in the mind of a participant about that model being in charge of controlling the stimulus, and instead provides support for one or more of the alternative models participants may be entertaining. This would result in participants increasing the probability that their choice will be guided by an alternative model and thus make a non-dominant choice. Similarly, when the previous choice of a participant is non-dominant, it reflects the fact that participants had low confidence that the dominant model was controlling the stimulus at the time and thus are likely to keep using whatever alternative model they entertain. We decided to focus uniquely on choices of the last task, because both the proportion of non-dominant stimuli and the proportion of non-dominant choices was the greatest of all tasks, thus allowing larger numbers of trials in the different groups. The stimulus sequence in this task was also common for all experiments (except for experiment 2).

### Logistic regression to confirm structure and gated learning

Looking at the proportion of dominant choices and how it depends on the trial history across all experiments provided strong evidence that both the error structure as well as the patterns of learning aligned well with the predictions from the MGM framework. However, the effects shown in those analyses are achieved by pulling average choice proportions across individuals. As a complementary analysis, we decided to run within participant regressions to predict trial-to-trial choices as a function of the recent history with the aim of confirming the structure in the errors as well as the gated learning mechanism. Thus, for each participant we ran a logistic regression model to predict their choice type (dominant choices coded as +1 and non-dominant ones as 0) in each trial of the fourth task, as a function of five predictors. The statistical inference was then made on the distribution of the resulting estimates, making the approach equivalent to a “mixed effects model”.

The first predictor signalled whether the previous choice was non-dominant or dominant (coded as +1 and −1, respectively, meaning that negative coefficients reflect a reduction in the probability of making a dominant choice). The second predictor signalled whether the latest stimulus was non-dominant or dominant (coded as +1 and −1, respectively, again meaning that negative coefficients reflect a reduction in the probability of making a dominant choice). The third predictor signaled the index of the trial in order to get a sense of whether performance was increasing through time (the index of the trial was z-transformed to have mean of zero and standard deviation of one but kept its ordinal nature). The last two predictors allowed us to assess the two facets of the gated learning mechanism by separating the effect of a three-way interaction between the three previous predictors into two simple effects (based on the previous choice type) that looked at the interaction between trial index and latest stimulus type.

Thus, the two last predictors (i.e., fourth and fifth predictors) both look at how the effect of non-dominant stimuli changes with time (interaction between time and latest stimulus type) but they separate the effects by the type of previous choice. The fourth predictor includes the crucial interaction term (trial index vs stimulus type) for trials when the previous choice was non-dominant (and has zeros elsewhere), thus reflecting the degree to which participants learn to deal with non-dominant stimulus transitions but limited to trials where the previous choice was non-dominant (i.e., when participants were unlikely to have been using the dominant model to guide their choices). Finally, the fifth predictor includes also the crucial interaction term (trial index vs stimulus type) but this for trials where the previous choice was dominant and zeros elsewhere. Mixture model learning predicts that participants will only update the parameters of their models (e.g., to incorporate more stochasticity) in proportion to the extent that each model is deemed to be responsible for the observation. Thus, we expect for the regression coefficient of the fifth predictor to be greater than the coefficient of the fourth predictor. To obtain reliable estimates in the regression, we made sure that we ran a regression only for participants that had at least six non-dominant choices out of all trials in the fourth task, this left 599 participants. One regression was run for each participant and statistical inferences were made on the distribution of regression coefficients over the participants by comparing the distribution against a null distribution around zero using a single-sample two-tailed t-test for the distribution of each coefficient.

## Supplementary materials

**Prediction and production task instructions**. Below, we present screenshots of the instructions given to participants at the start of the experiment.

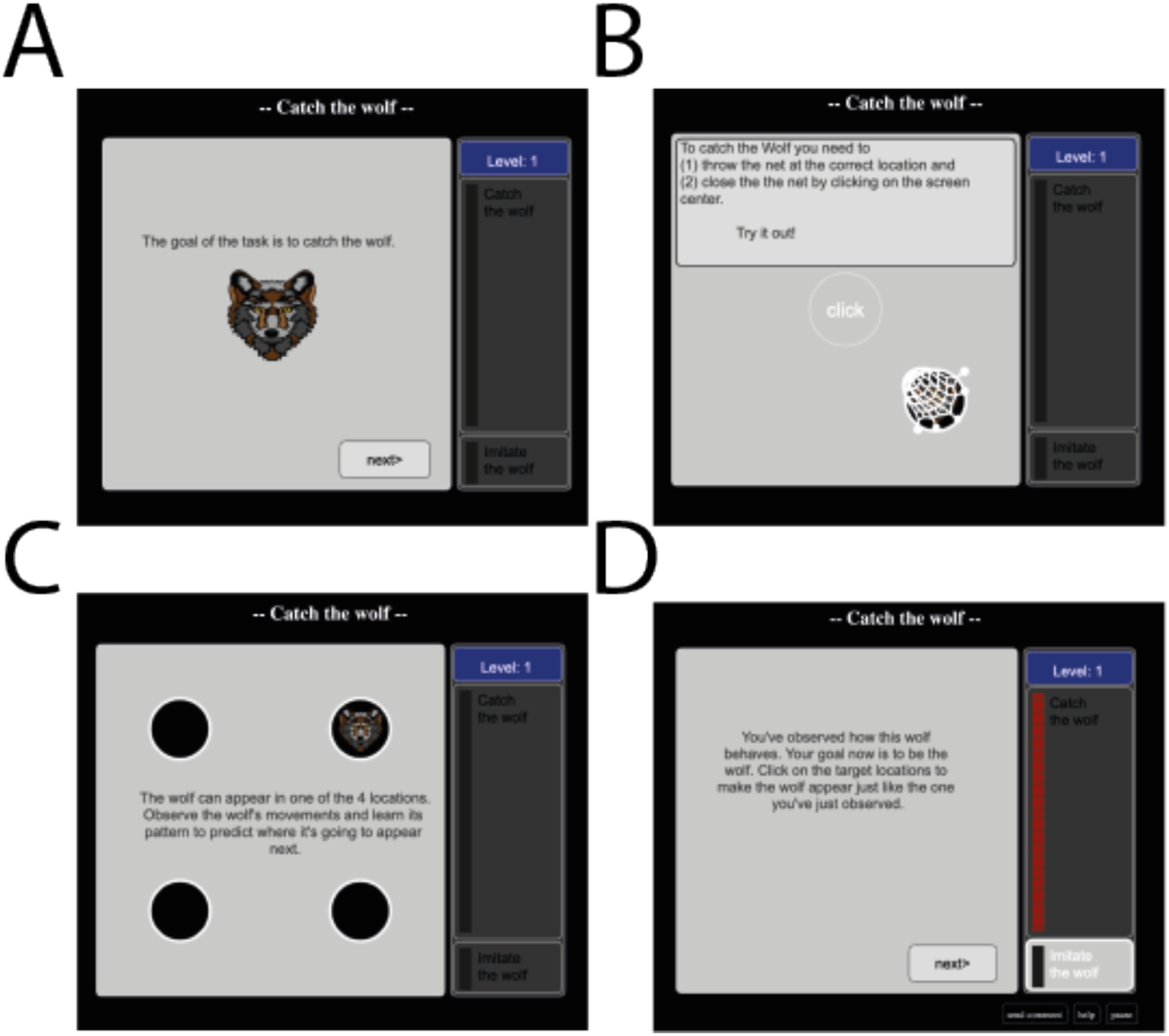

